# Graph convolutional network learning model based on new integrated data of the protein-protein interaction network and network pharmacology for the prediction of herbal medicines effective for the treatment of metabolic diseases

**DOI:** 10.1101/2024.04.24.591043

**Authors:** Tae-Hyoung Kim, Ga-Ram Yu, Dong-Woo Lim, Won-Hwan Park

**Author notes:** Corresponding Author (W.-H.P.), (D.-W.L.).

## Abstract

Chronic metabolic diseases constitute a group of conditions requiring long-term management and hold significant importance for national public health and medical care. Currently, in Korean medicine, there are no insurance-covered herbal prescriptions designated primarily for the treatment of metabolic diseases. Therefore, the objective of this study was to identify herbal prescriptions from the existing pool of insurance-covered options that could be effective in treating metabolic diseases. This research study employed a graph convolutional network learning model to analyze PPI network constructed from network pharmacology, aiming to identify suitable herbal prescriptions for various metabolic diseases, thus diverging from literature-based approaches based on classical indications. Additionally, the derived herbal medicine candidates were subjected to transfer learning on a model that binarily classified the marketed drugs into those currently used for metabolic diseases and those that are not for data-based verification. GCN, adept at capturing patterns within protein-protein interaction (PPI) networks, was utilized for classifying and learning the data. Moreover, gene scores related to the diseases were extracted from GeneCards and used as weights. The performance of the pre-trained model was validated through 5-fold cross-validation and bootstrapping with 100 iterations. Furthermore, to ascertain the superior performance of our proposed model, the number of layers was varied, and the performance of each was evaluated. Our proposed model structure achieved outstanding performance in classifying drugs, with an average precision of 96.68%, recall of 97.18%, and an F1 score of 96.74%. The trained model predicted that the most effective decoction would be *Jowiseunggi-tang* for hyperlipidemia, *Saengmaegsan* for hypertension, and *Kalkunhaeki-tang* for type 2 diabetes. This study is the first of its kind to integrate GCN with weighted PPI network data to classify herbal prescriptions by their potential for usage on certain diseases.

## 1. Introduction

In developed countries, the increase in the average lifespan has seen a concurrent rise in the prevalence of chronic metabolic diseases, including hyperlipidemia (HLP), hypertension (HTN), type 2 diabetes (T2D), and obesity [1]. These conditions are of growing importance to national public health and medical care, as they require continuous long-term management, thereby imposing significant financial burdens on national healthcare budgets [2]. The development of HLP, T2D, and HTN has been attributed to genetic predispositions, dietary habits, physical activity, and aging [3]. The World Health Organization (WHO) has identified HLP and HTN, along with alcohol consumption and smoking, as leading causes of death [4]. Additionally, the International Diabetes Federation (IDF) estimates that the number of people with diabetes reached 536.6 million in 2021 [5].

HLP is commonly managed through dietary modifications and the use of statin drugs, which inhibit the 3-hydroxy-3-methylglutaryl-coenzyme A (*HMG-CoA*) reductase enzyme, a key enzyme in the cholesterol synthesis pathway [6, 7]. HTN treatment primarily involves dietary adjustments such as weight loss, low sodium intake, increased potassium intake, and the use of antihypertensive medications, e.g. angiotensin receptor blockers and calcium channel blockers like amlodipine [8]. Additionally, for T2D, a wide range of oral hypoglycemic agents and insulin formulations are available for treatment of hyperglycemia over the past decades [9]. However, statin drugs, used to treat HLP, carry the risk of severe side effects, including myalgia, rhabdomyolysis and polyneuropathy [10]. Amlodipine, one of the leading agents for the treatment of hypertension, has side effects such as dizziness, flushing, and headaches [11], and in those patients with T2D who require daily insulin injections, treatment adherence is poor due to patient discomfort [9]. The development of new drugs requires an enormous expenditure of funds and time, with continually declining success rates [12]. Oriental medicine offers numerous herbal prescriptions with fewer side effects and excellent efficacy for patients requiring chronic disease management. However, in South Korea, among the 56 national insurance-covered herbal prescriptions available, there are currently none specifically designated for the treatment of metabolic diseases. Additionally, the development of new herbal medicines with insurance coverage is challenging due to a lack of institutional support, making it an objective difficult to achieve in the short term. Therefore, identifying effective herbal prescriptions for the treatment of metabolic diseases from the existing pool of insurance-covered herbal prescriptions and expanding their indications could be a feasible solution.

Herbal medicines are prepared by combining a variety of medicinal herbs, resulting in formulations that contain numerous bioactive compounds [13]. Therefore, a single prescription can be used for treating multiple diseases. The concept of drug repurposing can be applied to herbal medicine [14]; however, selecting the appropriate prescription is a critical step toward a successful outcome. Elucidating the efficacy of herbal medicines at the molecular level is a significant challenge due to the complexity that arises from the actions of multiple compounds present in these medicines on many targets. To analyze the intricate mechanisms and efficacy of herbal medicines, recent research has increasingly utilized network pharmacology and deep learning approaches.

Many studies utilize deep learning techniques to understand the complex target network data of pharmaceuticals and to categorize them [15–17]. Recently, researchers tried to apply deep learning to study the mechanisms of herbal prescriptions through methods such as natural language processing techniques based on ancient texts [18–21]. The graph convolutional network (GCN) is one of the deep learning models that can be specifically applied to protein-protein interaction (PPI) network data [22]. In addition, the method of applying protein weights for the GCN model based on the PPI network is different from the conventional organizing method of deep learning data based on the connectivity of the target network, and the gene scores of the PPI network genes can be sigmoid transformed and used as weights. Our data are the first to show how the graph data from PPI networks with assigned gene score values of drugs collected from Genecards can be trained as a GCN structure to deduce proper herbal prescriptions against certain disease [23] as weights to predict the effectiveness of a drug-specific protein combination for those diseases. We obtained data on the connectivity of the graphs in the target network of several approved drugs and herbal prescriptions whose current indications are metabolic diseases or non-metabolic diseases and utilized these as a test bed for the deep learning models.

We also formed a PPI network of existing drugs for metabolic diseases and applied the weights to the GCN model. The pre-trained model of drug data was trained for transfer learning on herbal prescription data [24]. As a result, we were able to classify drugs into those that are used for metabolic diseases and those that are not, which implied that our model could learn and classify data that are unique to drugs.

Before building our proposed model, we first obtained and organized large-scale multidisciplinary information on herbal prescriptions, including comprehensive data on herbal compounds, their targets, and disease-specific gene scores. To validate the generalization performance of the pre-trained model, we used metrics such as bootstrapping and K fold cross-validation after the classification of drugs for three representative metabolic disease groups: HLP, T2D, and HTN, currently in the market [25, 26]. Subsequently, to evaluate whether the herbal prescription data could be separated, we mixed positive and negative drug data together as references with the herbal prescription data and evaluated the performance by checking to see if they were classified correctly. The results showed that the pre-trained model separated the drugs used for diseases with good performance, and hence our study suggests that GCN learning using new data combinations can be used to recommend new herbal prescriptions for target diseases.

## 2. Materials and Method

### 2.1 Model Overview

To classify candidate herbal prescriptions for metabolic diseases, our proposed model was pre-trained based on GCNs with drugs whose main indication was one of the three metabolic diseases, HLP, T2D, or HTN. Subsequently, the generated model was transferred to the herbal prescription data to construct a dataset with the same structure to derive insured herbal prescriptions for each metabolic disease.

### 2.2 Data Preparation

We obtained drugs, herbal prescriptions, diseases, and PPI/compound-protein interaction (CPI) network data from the relevant online databases as described below. We processed and converted the data to make it suitable for GCN learning.

#### 2.2.1 Drugs

Drugs associated with metabolic diseases were identified from the statistical data of the Health Insurance Review and Assessment Service (HIRA, https://www.hira.or.kr, accessed on 24.01.15), and a binary classification was done, with drugs indicated for HLP, T2D, and HT assigned a positive value, and drugs used for other diseases assigned a negative value [27]. All data for training the pre-trained model are available in S1. Table.

#### 2.2.2 Herbal Prescription Database

The list of insured herbal prescriptions used in South Korea was collected from HIRA (https://www.hira.or.kr, accessed on 24.01.15). The constituent herbs and their content ratios in the 56 insured herbal prescriptions were obtained from the ‘Price List of Standard Prescription for Herbal Health Insurance provided by the National Health Insurance Service’. To construct a database of standard substance profiles of each insured formulation, our study collected and used all five major indicator substances provided for each herbal prescription from the pharmacological databases of natural products [28] of the Korea Food and Drug Administration (KFDA) (https://www.mfds.go.kr/, accessed on 24.01.15).

#### 2.2.3 Metabolic Disorder-related Targets

Gene scores for the protein related to HLP, T2D, and HT used in the study were collected from Genecards (https://www.genecards.org/, ver. 5.19), and the keywords used were Hyperlipidemia, Type 2 Diabetes, and Hypertension, respectively, and the search scope was limited to *Homo sapiens*. The collected data of gene list were 2,075 for hyperlipidemia, 17,916 for type 2 diabetes, and 11,637 for hypertension, respectively. The score values of the target genes associated with the diseases provided from Genecards were z-score normalized and sigmoid transformation was applied to represent the node importance within the PPI network before training [23]. In this study, the z-score normalization of the gene scores was converted according to the following formula:

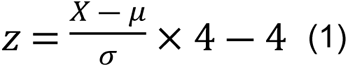

#### 2.2.4 Compound-Protein interaction (CPI), Protein-Protein interaction (PPI) Network

The connectivity information of the CPI network and drug-target score for each drug and core compound of the herbal medicine were collected through the search tool for interactions of chemicals (STITCH [ver. 5.0]), and the PPI network data of the target proteins regulated by each drug were collected by transferring the gene list obtained from STITCH into the Search Tool for the Retrieval of Interacting Genes/Proteins (STRING [ver. 12.0]). The PPI network data collected for each drug was then classified into binary values according to whether there was a connection between each of the proteins and used as raw data for the GCN training [29, 30].

### 2.3 Deep Learning Approach

In this research study, we used the GCN model for deep learning based on the theory of network pharmacology. This is the first study that has used the PPI and CPI networks of herbal prescriptions as a list of the controllable target proteins of compounds and their connections to each other, and the gene scores of the Genecards as weights for the relevant proteins of each network component to indicate the importance of the protein in HLP, T2D, and HTN. The parameters of the model for pre-learning can be found in S2. Table.

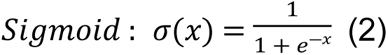

GCN is a model that is specialized for learning complex graph structures, therefore, the setting is applicable to this study. GCNs work by updating attributes connected through nodes and edges inside a graph [31].

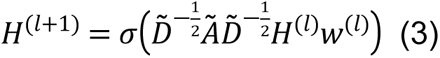

As there are only a few insurance-covered herbal prescriptions and none of them are currently used for metabolic diseases as the main indication, a model was trained by classifying the drugs into those currently used for metabolic diseases and drugs used for other indications. The generated model was transferred to herbal medicines and designed to classify herbal prescriptions in a manner identical to drugs rather than herbal prescriptions, independently based on classical indications from literature or empirical evidence from oriental medicine practitioners.

We checked the constituent compounds of herbal prescriptions and their target proteins from the database and transformed them into the CPI network, and then into the PPI network.

### 2.4 Performance Validation of the Deep Learning Model

Our proposed model was the first target weight-based learning model for drug-based networks and there was no comparable previous work. Therefore, we needed to validate the generalization ability of the model and ensure that it did not overfit the training data. We used the most widely used K-fold cross-validation and bootstrapping techniques to evaluate the reliability of the model in predicting herbal prescriptions[25, 26]. All analyses were carried out using Python3.11.6 [32], and Torch 2.1.0 [33] with Visual Studio Code 1.85.1.

#### 2.4.1 5-Fold Cross Validation

5-fold cross-validation divides the entire dataset into five parts and uses one of them as the validation data for all the data and repeats it five times for the remaining data as training data. This method allows the validation of the results on all datasets and the evaluation of the generalizability of the performance [26]. We trained the model using marketed drug data and used it in this study.

#### 2.4.2 Bootstrapping

Bootstrapping regenerates bootstrap samples by random sampling from the dataset and using the newly generated sample dataset to learn with as many iterations as you set. We used the method to train the model 100 times to evaluate its generalization ability [25].

#### 2.4.3 Confusion Matrix

The Confusion Matrix is used to evaluate the performance of a model in a classification problem [34]. The data predicted by the model is divided into TP, TN, FP, and FN as below, to check the learning results, and the data can be used to calculate performance metrics such as accuracy, precision, recall, and the F1 score [35].

● ***True Positive (TP): Actual label is positive, and the model correctly predicted it as positive*.**
● ***True Negative (TN): Actual label is negative, and the model correctly predicted it as negative*.**
● ***False Positive (FP): Actual label is negative, but the model predicted it as positive*.**
● ***False Negative (FN): Actual label is positive, but the model predicted it as negative*.**
● 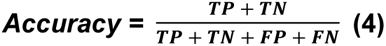
● 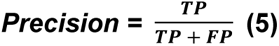
● 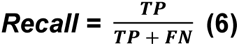
● 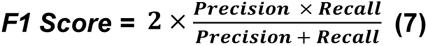

#### 2.4.5. Classifying the Herbal Prescriptions by New Indications Using the Trained Model

We aimed to verify whether the model could properly classify herbal prescriptions as potential drugs for diseases and evaluated the model by adding some drug data unused for HPL model training, as references. Therefore, we added actual marketed drugs used to treat metabolic diseases to verify that our model could classify real drugs and thus added reliability to the selected herbal prescription results.

## 3. Results

To build the pre-training model, we set the drug data corresponding to each disease among the 59 drugs as positive values and the other data as negative values. After pre-training, we derived the herbal prescriptions that were predicted to be available for each disease by transfer learning. The results of the generalization performance of the proposed model are shown, and the results of predicting herbal prescriptions based on the evaluation of the proposed model are presented.

### 3.1 Confusion Matrix of Pre-training Data

One method of evaluating a model’s performance is the Confusion Matrix, which is one of the popular generalized performance metrics and is explained in Fig 3. In our proposed model, 19 out of 22 HLP drugs were correctly classified, while 25 and 29 out of 32 T2D and HTN drugs, respectively, were correctly classified. The classification performance of the HLP classification model was 86.36%, that of the T2D classification model was 78.12%, and that of the HTN model was 90.06%.

**Fig. 1.**
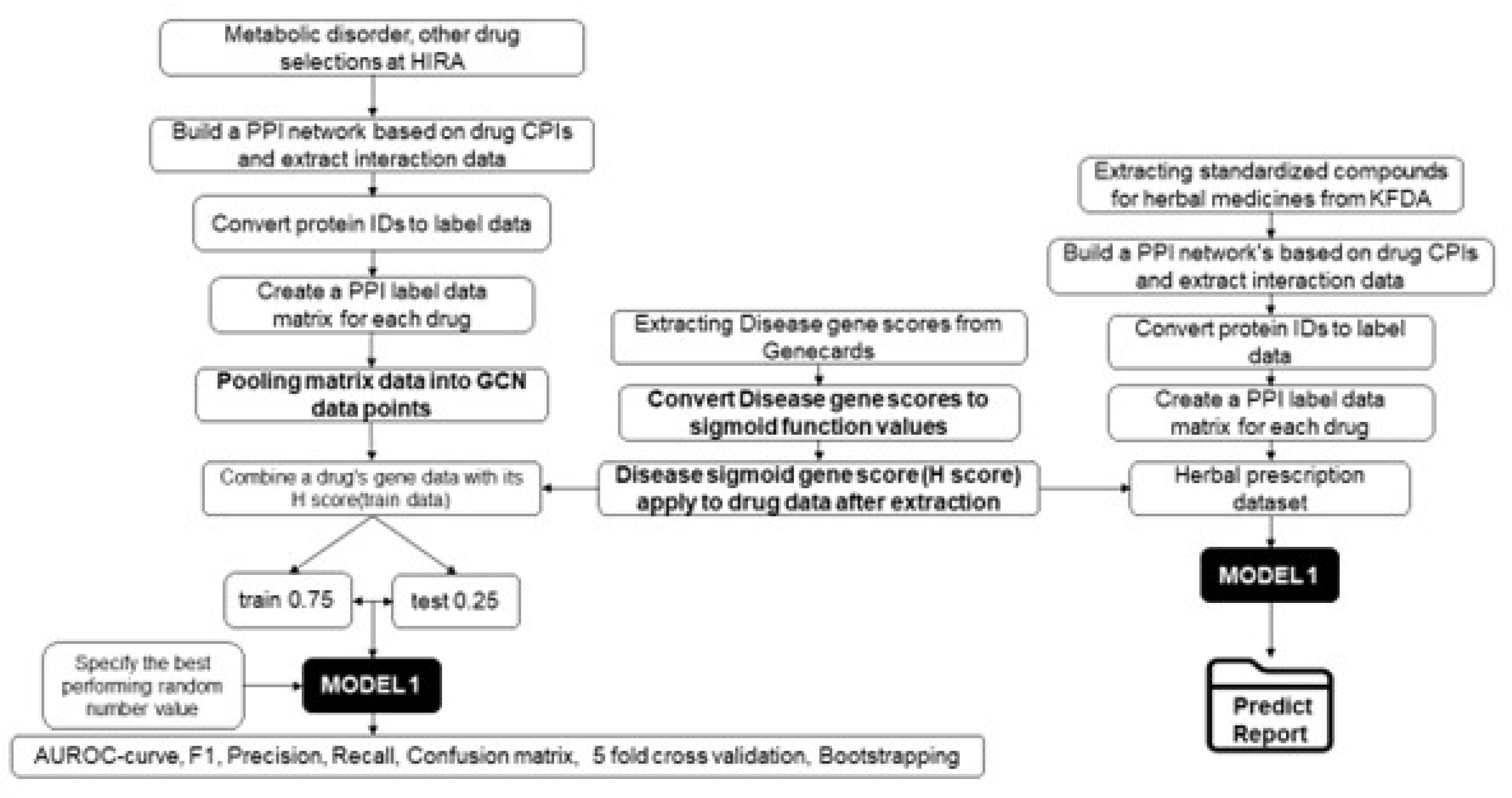
A complete framework for deriving herbal prescriptions for metabolic diseases by pre-training the model on marketed drug data and subsequently applying it to herbal prescription data to derive herbal prescriptions for metabolic diseases.

**Fig. 2.**
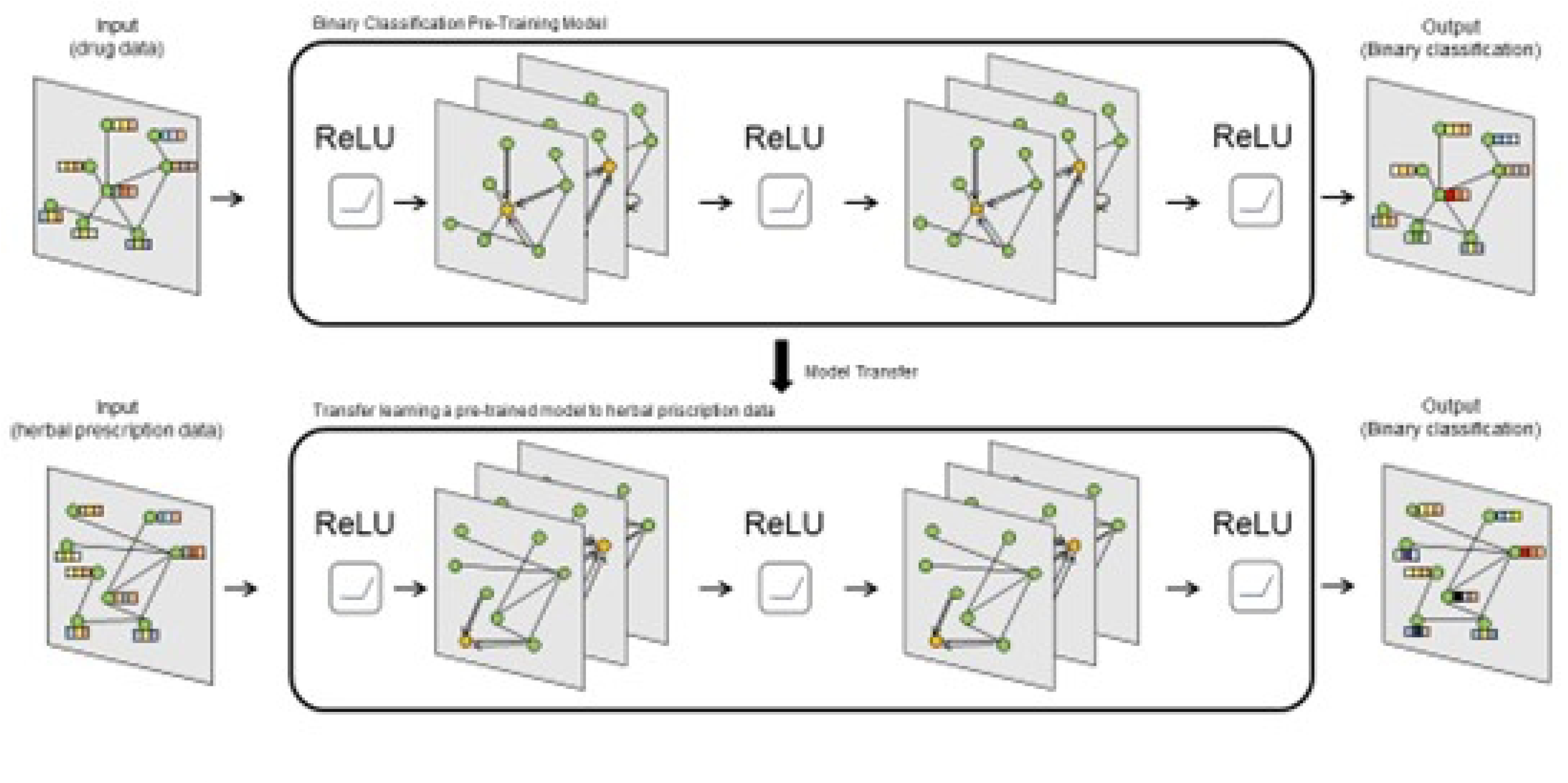
**Schematic of transfer learning to identify effective herbal prescriptions for metabolic diseases. After pre-learning, the binary classification of drugs into those used for metabolic diseases and those not used for metabolic diseases was done and the generated model was applied to herbal prescriptions to derive effective herbal prescriptions for metabolic diseases.**

**Fig 3.**
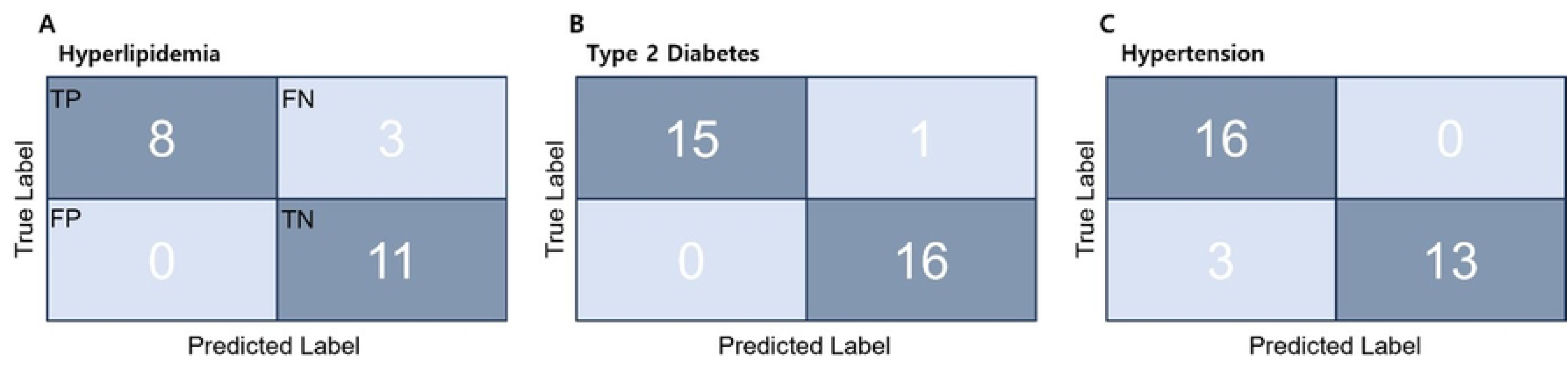
**The Confusion Matrix shows how well the model performed on the entire training data. A) Confusion Matrix for hyperlipidemia, B) Confusion Matrix for type 2 diabetes, and C) Confusion Matrix for hypertension.**

### 3.2 Bootstrapping to Identify the Best-performing Model Structure

To evaluate the performance of the model structures, we pre-trained the model with different layer depths and bootstrapped the results to get the average after 100 iterations for each layer. Fig 4 shows that our proposed model had the most stable performance compared to the other layer structures.

**Fig. 4.**
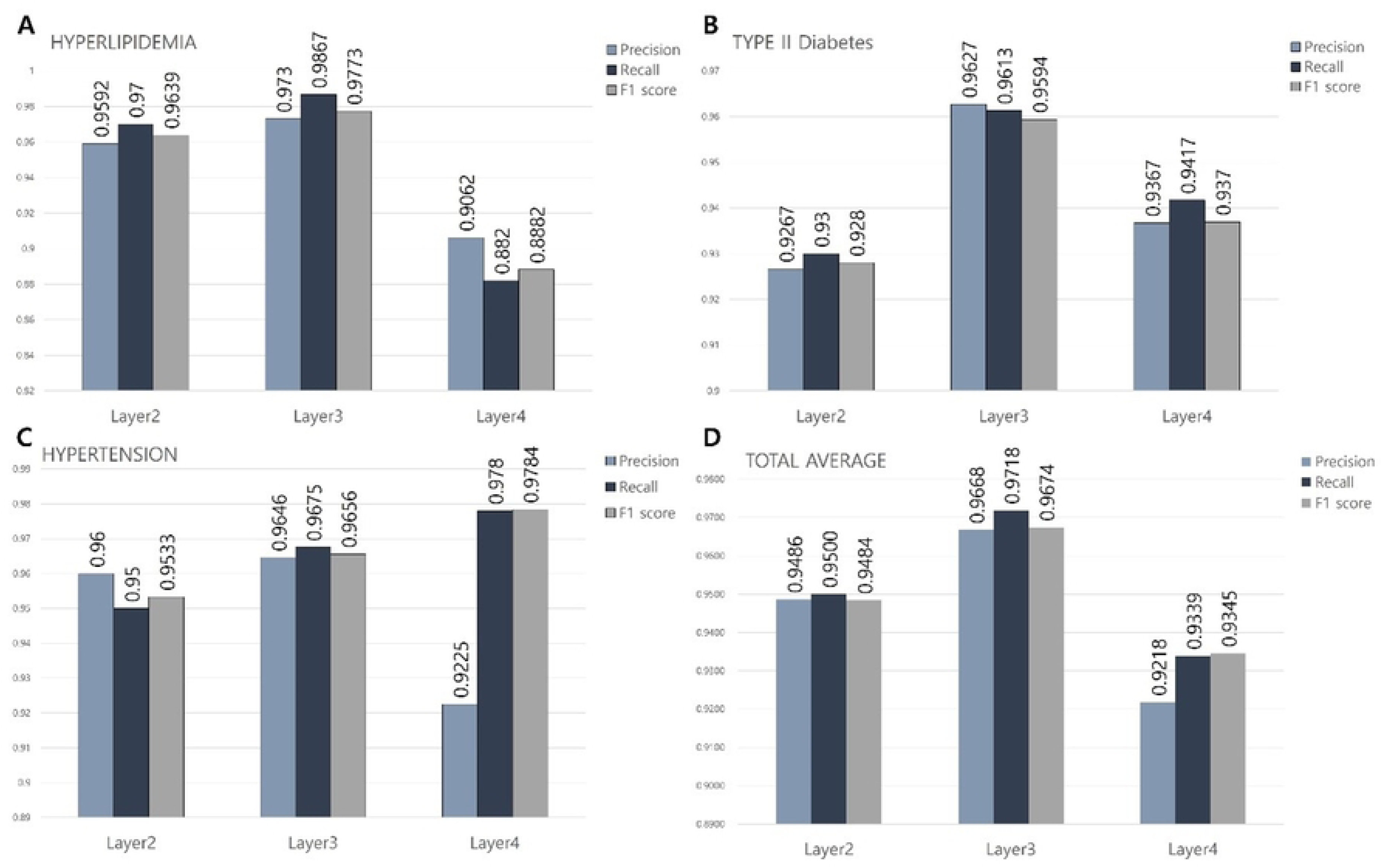
**Comparison of the validation metrics of the proposed graph convolutional network (GCN) model according to the layer depth to find the optimal model structure. A) Results by layer for Hyperlipidemia. B) Layer-by-layer results for Type 2 Diabetes C) Layer-by-layer results for Hypertension D) Average of the results of the three disease models.**

### 3.3 5-Fold Cross-Validation

The results of 5-fold cross-validation for the model are presented in Table 1. It showed that all datasets had a high learning rate and low loss on the validation data, indicating that the model had good generalization ability. The average loss value of the 5-fold cross-validation results for this model was 0.3232 and the average learning rate was 0.8501.

**Table 1.** Loss and learning rate were verified by 5 fold cross validation for HLP, T2D and HTN respectively.

|  | Hyperlipidemia |  | Type 2 Diabetes |  | Hypertension |  |
| --- | --- | --- | --- | --- | --- | --- |
|  | Accuracy | Loss | Accuracy | Loss | Accuracy | Loss |
| Fold 1 | 0.8000 | 0.4908 | 1.0000 | 0.2339 | 0.7143 | 0.4245 |
| Fold 2 | 1.0000 | 0.1213 | 1.0000 | 0.2821 | 1.0000 | 0.2066 |
| Fold 3 | 1.0000 | 0.2343 | 1.0000 | 0.1262 | 0.8333 | 0.3529 |
| Fold 4 | 0.7500 | 0.4877 | 1.0000 | 0.1348 | 1.0000 | 0.2789 |
| Fold 5 | 0.7500 | 0.4529 | 0.8333 | 0.3060 | 1.0000 | 0.1474 |
| Average | 0.8600 | 0.3105 | 0.9667 | 0.2166 | 0.9095 | 0.2821 |

### 3.4 Results of Classification of Herbal Prescriptions by Diseases

To verify the generalization of the pre-trained model and to apply the model to herbal prescriptions, we collected all available candidate herbal prescription data for metabolic diseases (Table 2). The results of all the herbal prescriptions are presented in S3. Table.

**Table. 2.** Top 10 herbal prescriptions predicted by the pre-trained model to be effective for hyperlipidemia, type 2 diabetes, and hypertension, respectively, after analyzing the entire herbal prescription data. The lower the score, the more effective the herb is predicted to be for that condition.

| Hyperlipidemia |  | Type 2 Diabetes |  | Hypertension |  |
| --- | --- | --- | --- | --- | --- |
| Name | Score | Name | Score | Name | Score |
| Jowiseunggi-tang | 0.377 | Kalkunhaeki-tang | 0.214 | Saengmaegsan | 0.175 |
| Dangkwiughwangtang | 0.392 | Kungso-san | 0.250 | Banhahubaktang | 0.420 |
| Samhwangsasim-tang | 0.395 | Naesosan | 0.283 | Jowiseunggi-tang | 0.425 |
| Injinhotang | 0.415 | Danggwi Yeongyoeum | 0.316 | Naesosan | 0.453 |
| Gu-Mi-Gang-Hwal-Tang | 0.423 | Hyeonggaeyeongyotang | 0.340 | Injinhotang | 0.48 |
| Hoechunyanggyeok-san | 0.424 | Bokryungbosim-tang | 0.347 | Leejung-tang | 0.488 |
| Pyeong wi san | 0.438 | Ohjuksan | 0.384 | Gamisoyosan | 0.494 |
| Sosiho-tang | 0.458 | Bokryungbosim-tang | 0.383 | Pyeong wi san | 0.526 |
| Sihocheonggan-tang | 0.466 | Gu-Mi-Gang-Hwal-Tang | 0.428 | Bakchul-tang | 0.530 |
| Banhasasim-tang | 0.467 | Galguntang | 0.445 | Palmul-tang | 0.581 |

## 4. Discussion

Herbal prescriptions include two or more herbs, and their extracts contain numerous bioactive compounds. This makes it difficult to understand or predict the efficacy and targets of herbal medicines in the same way as synthetic drugs with a single target. In addition, the lack of standardized research on the components of herbal prescriptions and uniformity in efficacy/disease interpretation of the herbal prescription compared to traditional medical concepts or theories and modern medicine makes it difficult to explain the efficacy of herbal medicines to patients. To move forward as an evidence-based medicine (EBM), it is necessary to scientifically examine and re-evaluate the effectiveness of herbal prescriptions with objective data and analyses rather than empirical theories. The unique and complex substance profiles of many different herbal prescriptions have long posed a challenge to the study of their efficacy and mechanisms. Many studies have attempted to establish an objective basis for the efficacy and mechanisms of herbal prescriptions through network pharmacology, which is one of the methods to provide evidence based on the modern pharmacological interpretation of herbal prescriptions. Network pharmacology, based on the understanding of system biology, has been used to interpret various data obtained through the Kyoto Encyclopedia of Genes and Genomes (KEGG) pathway, Gene Ontology (GO), and the PPI network to reveal the mechanisms of herbal prescriptions to regulate various diseases in contrast to the past when such a study was not possible [36–38]. In addition, the exploration of new indications and diseases for drugs based on this network pharmacology, or drug discovery, has had some success [38–40]. However, these network pharmacology and PPI network analyses have the limitation that they are open to various interpretations, and the role of the final interpretation and selection of the result has been left to the researcher, as the data presented does not provide a selection or relative ranking of herbal prescriptions. Specifically, in studies using current clinical prescriptions, it is often difficult for researchers to make objective decisions because they have subjective expectations and biases regarding the efficacy of the prescriptions and the indications for which they have been used. In research using herbal prescriptions, it is necessary to overcome this subjective bias of the researcher and decide the efficacy for areas and indications that have not yet been identified based on objective criteria.

GCNs have been used in many fields of medicine to understand and predict the pharmacological mechanisms of herbal prescriptions and over-the-counter drugs. In a previous study with a purpose similar to ours, Zhao et al. constructed a network using Drug Target Interaction (DTI) based on the GCN model for drug development, extracted each feature to identify the mechanism of the drug through learning, and predicted the interaction of ligand-protein, protein-protein, etc. to discover new drug candidates [41]. Liu used machine learning to construct a heterogeneous network on all formula data to understand traditional herbal prescriptions and interpret the herb node embeddings to predict formulation efficacy and when compared to the results of other types of learning models, GCN learning was observed to be more effective than typical machine learning methods [42]. As such, GCNs have been studied in the field of pharmacology to understand drug mechanisms.

The study’s exploration of new indications aimed to break away from the so-called ‘Oriental medicine theory and experience-based biases’ with respect to conventional (old) indications and targets of prescriptions. Our study fused information from various natural product/drug and disease databases with connectivity data from the PPI network, which is the main basis of network pharmacology, and reinterpreted the connectivity graphs of the targets per prescription as pharmacological activity fingerprints. After training a model using deep learning with these pharmacological activity fingerprints as the input data, we present a new research method that deduces the potential of using a specific prescription for the presented disease.

Since there is no previous research using the same combination of data as ours, we rigorously checked the appropriateness of our proposed model. We evaluated which model performed the best by layer factor, pre-trained over-the-counter medicines with that model, and classified herbal prescriptions through transfer learning. As a result, we found that the model for HLP showed the highest probability of using *Jowiseunggi-tang*, followed by *Dangkwiyughwangtang* and *Samhwangsasim-tang.* Among them, *Jowiseunggi-tang,* the most probable candidate, is actually a prescription used in clinics for indications such as indigestion, constipation, and abdominal pain, but is expected to exert efficacy on many metabolic diseases [43–45]. As support to this opinion, serum experiments in an obese mouse model demonstrated that *Jowiseunggi-tang* significantly increased high-density lipoprotein cholesterol (HDL-C) and decreased total cholesterol, low-density lipoprotein cholesterol (LDL-C), and triglycerides when compared to the high-fat diet obese group [46]. Also, *Kalkunhaeki-tang*, which is considered useful in diabetes, has been commonly used for colds and headaches according to original indications from the medical classics [47]. However, no studies of its use in diabetes have been conducted yet, which could be an interesting topic for further research. Finally, *Saengmaegsan*, which was predicted to be used for HTN, has been used for heart disease, including arrhythmias and heart failure [48]. The study result suggests that *Saengmaegsan* may have a significant effect on HTN by reducing obesity rates, one of the main causes of hypertension [49]. These studies add credibility to the outcomes of our proposed GCN model [46, 49].

Thus, our results present a new interpretation of the role of herbal prescriptions, distinct from the old classic medical indications. The combination of the proposed GCN deep learning method, network pharmacological approaches, and data processes of the drug/herbal prescription target network connectivity data may contribute to further the development of oriental medicine and provide patients with a wider range of treatment options by presenting new herbal prescriptions or drugs. Our research categorized 59 drugs into treatments for four groups (HLP, T2D, HTN and none of these diseases as negative) used three models to predict the available herbal prescriptions for each condition. However, these data are not enough for deep learning training, and more drug data is needed to improve the performance of the model. Although this research deduced herbal prescriptions through computing science, it is necessary to verify the efficacy of the herbal prescriptions through in vivo experiments to use them in clinical practice. In addition, the study was based on the indicator substances for each herbal prescription provided by the KFDA, but in reality, there are many more compounds in each herbal prescription. It is necessary to further reinforce the data for each herbal prescription to use it in clinical practice.

## 5. Conclusion

To identify new indications of herbal prescriptions for metabolic diseases, we pre-trained a GCN deep learning model with data of drugs currently available in the market comprising their PPI network and weight scores according to the targeted metabolic disease. The performance of the model was validated through 5-fold cross-validation and bootstrapping with each of the three diseases, and the binary classification result showed successful differentiation of marketed drugs as per their indications. The transferred model was applied to the classification of herbal prescriptions with their PPI network data. The model predicted that among the 56 insurance-covered herbal prescriptions, HLP could be treated with *Jowiseunggi-tang*, T2D with *Kalkunhaeki-tang*, and HT with *Saengmaegsan*.

## Institutional Review Board Statement

Not applicable.

## Informed Consent Statement

Not applicable.

## Abbreviations

HLP: Hyperlipidemia
T2D: Type 2 Diabetes
HTN: Hypertension
GCN: Graph Convolutional Network
PPI: Protein-Protein Interaction
WHO: World Health Organization
HMG-CoA: Hydroxy-Methylglutaryl-coenzyme A
CPI: Compound-Protein Interaction
KFDA: Korean Food and Drug Administration
HIRA: HEALTH INSURANCE REVIEW & ASSESSMENT SERVICE
DTI: Drug-Target interaction
GO: Gene Ontology
KEGG: Kyoto Encyclopedia of Genes and Genomes

## Acknowledgments

Not applicable. All relevant data are within the manuscript and its Supporting Information files.

## CRediT authorship contribution statement

Tae Hyoung Kim: Methodology, Investigation, Original draft writing;

Ga-Ram Yu: Writing – review & editing;

Dong Woo Lim: Conceptualization, Supervision, Original draft writing;

Won-Hwan Park: Supervision, Funding

## Data Availability Statement

All data that support the findings of this study are included.

## Code availability

The code used in this study has been made open-source, available on GitHub https://github.com/Rapoudok/GCN-herbal-medicine

## Supporting information

S1 Table. Drug list. The classification result of a pre-trained model of metabolic diseases. Positive are drugs that are used for the disease and negative are drugs that are used for other diseases.

S2 Table. Parameters types. The types of parameters used in the pre-trained model for metabolic diseases.

S3 Table. Full result for herbal. is the result of applying all 56 insurance herbal prescriptions to the pre-trained model. 0 is the herbal prescription predicted to be available for the target disease and 1 is the herbal prescription predicted not to be available for the disease.

